# Mapping the GCK–GKRP interaction landscape using deep mutational scanning by reverse two-hybrid screening

**DOI:** 10.64898/2026.01.15.699699

**Authors:** Sarah Gersing, Sofie Ø. Dahl, Kresten Lindorff-Larsen, Rasmus Hartmann-Petersen

## Abstract

Glucokinase (GCK) regulates insulin secretion in the pancreas and glycogen synthesis in the liver. The best characterized interaction partner of GCK is the glucokinase regulatory protein (GKRP). GKRP is expressed in the liver where it regulates GCK activity and abundance. In this study, we used a library of GCK single amino acid substitutions and a reverse two-hybrid screen to characterize the impact of GCK variants on GKRP interaction. We obtained interaction scores for 7534 GCK single amino acid substitutions. A major driver of loss of interaction appeared to be loss of protein abundance, and we therefore compared interaction scores with our previously obtained GCK abundance scores to disentangle variant effects on interaction and abundance. Variants specifically perturbing interactions were found at the interaction interface but also structurally far from GKRP, revealing potential allosteric effects. In addition, unstable variants at buried residues in the large domain were stabilized by GKRP. These variants could potentially also be stabilized by e.g. pharmacological chaperones to rescue GCK function. In conclusion, we have now characterized the activity, abundance and interaction of 7128 GCK variants. Collectively these scores help pinpoint variants solely perturbing GKRP interaction, giving rise to liver-specific effects and potentially subtle glucose homeostasis phenotypes.

## Background

Appropriate glucokinase (GCK) function is crucial for glucose homeostasis, emphasized by variants in the *GCK* gene, which are linked to three glucose-homeostasis diseases [1]. GCK maintains glucose homeostasis by regulating insulin secretion in pancreatic beta cells [2, 3, 4] and glycogen synthesis in hepatocytes [5, 6, 7]. In these two pathways, the rate limiting step is the phosphorylation of glucose to form glucose 6-phosphate, a reaction catalyzed by GCK [5, 6]. The activity of GCK thus determines glycolytic flux, and is optimized to maintain glucose homeostasis. Specifically, GCK has a low glucose affinity (*S*_0.5_ 7.5–8.5 mM) and a sigmoidal kinetic response to changes in glucose levels (*n*_*H*_ 1.7) [8, 9, 10], enabling GCK to maintain appropriate blood glucose levels. Variants that alter GCK function are linked to three glucose-homeostasis diseases: permanent neonatal diabetes mellitus (PNDM) (MIM# 606176), GCK-maturity-onset diabetes of the young (GCK-MODY) (MIM# 125851) and hyperinsulinemic hypoglycemia (HH) (MIM# 601820). Variants that increase GCK activity are linked to HH, while variants that decrease activity are linked to PNDM when homozygous or compound heterozygous, and to GCK-MODY when heterozygous. Diagnosis of *GCK*-linked diseases relies on DNA sequencing and interpreting the effect of any identified variants.

To facilitate variant interpretation, we have previously characterized the activity and abundance of nearly all (97% and 95%, respectively) single amino acid substitutions in GCK [11, 12]. By broadly assaying GCK variant activity, we identified both hypo- and hyperactive variants to facilitate diagnosis of GCK-linked diseases. We next examined GCK variant abundance, as decreased protein stability is one of the dominating mechanisms for loss-of-function variants [13, 14]. Accordingly, 43% of GCK single amino acid substitutions potentially decrease activity through a reduced protein abundance.

One important aspect of GCK function, which has not been characterized in depth, is the regulation of GCK by glucokinase regulatory protein (GKRP). GKRP is expressed in the liver where it inhibits GCK activity [15]. When GKRP binds GCK, it prevents substrate binding, and thereby GKRP acts as a competitive inhibitor with respect to glucose. At low glucose levels, GKRP binds and sequesters GCK in the nucleus [16, 17], while at higher glucose levels the complex dissociates to enable GCK-mediated glycogen synthesis in the cytosol. When bound to GKRP, the GCK conformation closely resembles the inactive (super-open) state which dominates at low glucose levels [18], as seen from the structure of the GCK–GKRP complex [19]. The crystal structure of the complex was solved using rat liver GKRP and the pancreatic isoform of human GCK [19]. As GKRP is not expressed in the pancreas, only the hepatic GCK isoform interacts with GKRP in vivo. However, the two isoforms are encoded by the same gene and only differ in the 15 N-terminal residues [20], which are not involved in the GKRP interaction. The GCK–GKRP interaction, in addition to inhibiting GCK, also appears to stabilize GCK, as loss of GKRP leads to reduced GCK protein levels [17, 21]. Potentially, the dual role of GKRP in GCK activity and stability might explain the relative mild effects observed in mice without GKRP [17, 21].

The importance of GKRP for appropriate GCK activity has, however, been underscored by the common GKRP allele P446L (minor allele frequency 34% [22]), which is associated with raised triglyceride levels and reduced fasting blood glucose [22]. The GKRP P446L variant shows decreased interaction with GCK [23] and also confers decreased GCK protein abundance [24]. Conversely, the effects of GCK variants on GKRP interaction have also been studied for some disease-causing GCK variants, but also more broadly using a novel reverse two-hybrid system [25]. In that study, variants perturbing GCK–GKRP interaction were primarily found at or near the interaction interface. However, only 12 single amino acid substitutions with loss of interaction were examined.

To more broadly characterize the effects of GCK variants on GKRP interaction, we here used a reverse two-hybrid screen to characterize the GCK–GKRP interaction for 7534 GCK amino acid substitutions. We found that variants showing reduced GKRP interaction concentrated both at the interaction interface, as expected, and more generally in buried residues, likely due to indirect (protein stability) effects. By comparing interaction scores with our previously obtained GCK abundance scores [12], we identified variants that specifically perturbed GKRP interaction, but also low abundance variants that were stabilized by GKRP. Potentially, these variants could be therapeutically targeted with e.g. pharmacological chaperones to stabilize the GCK protein and rescue function.

## Results and Discussion

### Variant impact on the GCK–GKRP interaction using a reverse two-hybrid system

To examine the impact of GCK variants on GCK–GKRP interaction, we used the reverse two-hybrid system. Here, GCK (the prey) is expressed as a fusion protein with the GAL4 activation domain (GAL4-AD), and GKRP (the bait) is expressed as a fusion protein with the GAL4 DNA binding domain (GAL4-DBD). The two fusion proteins are expressed in a yeast strain (MaV203) with genome-integrated reporter genes, including *URA3*. Specifically, the regulatory region of the *URA3* gene contains the DNA binding sites for the GAL4-DBD. Accordingly, if GCK and GKRP interact, this brings the two GAL4 domains together, leading to expression of *URA3*. GCK variants that interact with GKRP will therefore enable yeast growth on medium without uracil. Conversely, interacting variants will grow poorly on medium containing 5FOA, which kills yeast cells expressing *URA3*, and we could therefore select for GCK variants that decreased GKRP interaction by growing yeast on 5FOA medium (Fig. 1A).

**Figure 1.**
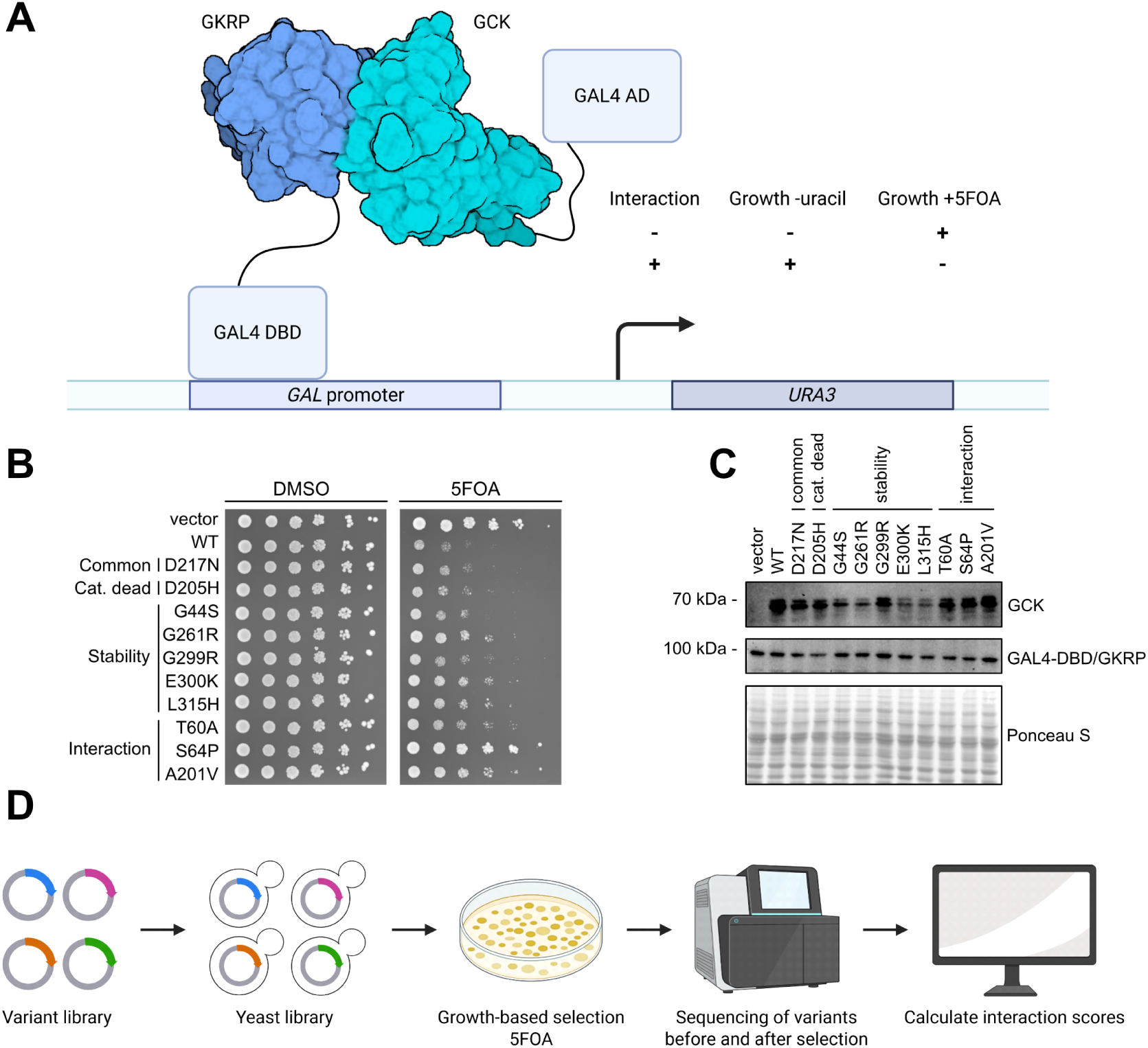
Measuring GCK–GKRP interaction using a reverse two-hybrid assay. (A) Illustration of the reverse two-hybrid assay. In yeast, GCK is expressed fused to the GAL activation domain (GAL4 AD) and GKRP is expressed fused to the GAL DNA-binding domain (GAL4 DBD). When GCK and GKRP interact, GAL4 drives the expression of reporter genes including *URA3*, enabling the cell to grow without uracil, while inhibiting growth in presence of 5FOA. Created in BioRender. Hartmann-Petersen, R. (2026) https://BioRender.com/yfpa1dw. (B) Small scale reverse two-hybrid growth assay with yeast co-expressing GAL4-DBD-GKRP and GAL4-AD-GCK wild type and the shown variants. As a control, cells were grown on DMSO, while 5FOA selected for variants with decreased interaction. (C) Western blot showing the protein levels of wild-type GCK and variants in the reverse-two hybrid setup. (D) Illustration of the multiplexed reverse two-hybrid assay. Created in BioRender. Hartmann-Petersen, R. (2026) https://BioRender.com/5vfgi9i.

To test the reverse two-hybrid system for GCK–GKRP, we co-expressed GAL4-DBD-GKRP with wild-type GAL4-AD-GCK or ten GCK missense variants (Fig. 1B). These test variants included the benign variant D217N [26], expected to behave similar to wild-type, the catalytically dead variant D205H [18, 27], also not expected to affect interaction, five low-abundance variants (G44S, G261R, G299R, E300K, L315H) [12], and three variants previously shown to perturb GCK–GKRP interaction (T60A, S64P, A201V) [25]. When grown on 5FOA medium, wild-type GCK grew poorly compared to a pDEST22 vector control, showing that wild-type GCK interacts with GKRP in this system. As expected, D217N and D205H also showed poor growth, again suggesting a strong interaction with GKRP. Similarly, the low-abundance variant G44S also showed wild-type-like interaction. In contrast, the seven remaining variants showed various levels of increased growth, suggesting that these variants perturb interaction. When we compared the reverse two-hybrid assay with a western blot showing the protein levels of wild-type GCK and the ten variants (Fig. 1C), the five variants, known to affect GCK stability, as expected, decreased GCK protein levels, and therefore likely decreased GCK–GKRP interaction through decreased protein abundance. The three variants, previously shown to perturb interaction, decreased interaction specifically, as they did not perturb protein abundance. In conclusion, the reverse two-hybrid system can assay GCK–GKRP interaction, with loss of abundance or direct perturbation of the interaction being two possible sources for decreased interactions.

Having established a growth assay to probe GCK–GKRP interaction, we next multiplexed the assay to widely assess the interaction of GCK variants with GKRP (Fig. 1D). Given that we already had a variant library for the pancreatic GCK isoform, we used this library to characterize the GCK–GKRP interaction. For the reverse two-hybrid screen, we first cloned the library of GCK variants ([11]) into the pDEST22 destination vector, encoding the GAL4-AD. This library was then transformed into the reverse two-hybrid yeast strain already expressing GAL4-DBD-GKRP. The reverse two-hybrid library was grown on 5FOA medium to select for variants with decreased GKRP interaction. Then, plasmids were extracted from yeast before and after selection and the *GCK* ORF was amplified in tiles [28]. These amplicons were sequenced and analyzed to determine the change in frequency of variants following 5FOA selection and thereby their impact on GCK–GKRP interaction. In this way, we obtained interaction scores for 7534 variants (81.2% of the possible single amino acid variants). An interaction score of 0 indicates interaction similar to nonsense variants, while an interaction score of 1 indicates interaction similar to synonymous (wild-type-like) variants.

### A map of GCK–GKRP interaction

The heatmap of GCK variant impact on GKRP interaction showed that variants decreasing interaction are spread throughout the GCK sequence (Fig. 2A). As expected, nonsense variants severely decreased interaction except in the C-terminal region, which appeared to be dispensable for interaction. As expected, residues where variants severely decreased interaction were often found at and near the interaction interface. In addition, many residues distant in sequence to interacting residues were also important for interaction, potentially indirectly by affecting protein stability.

**Figure 2.**
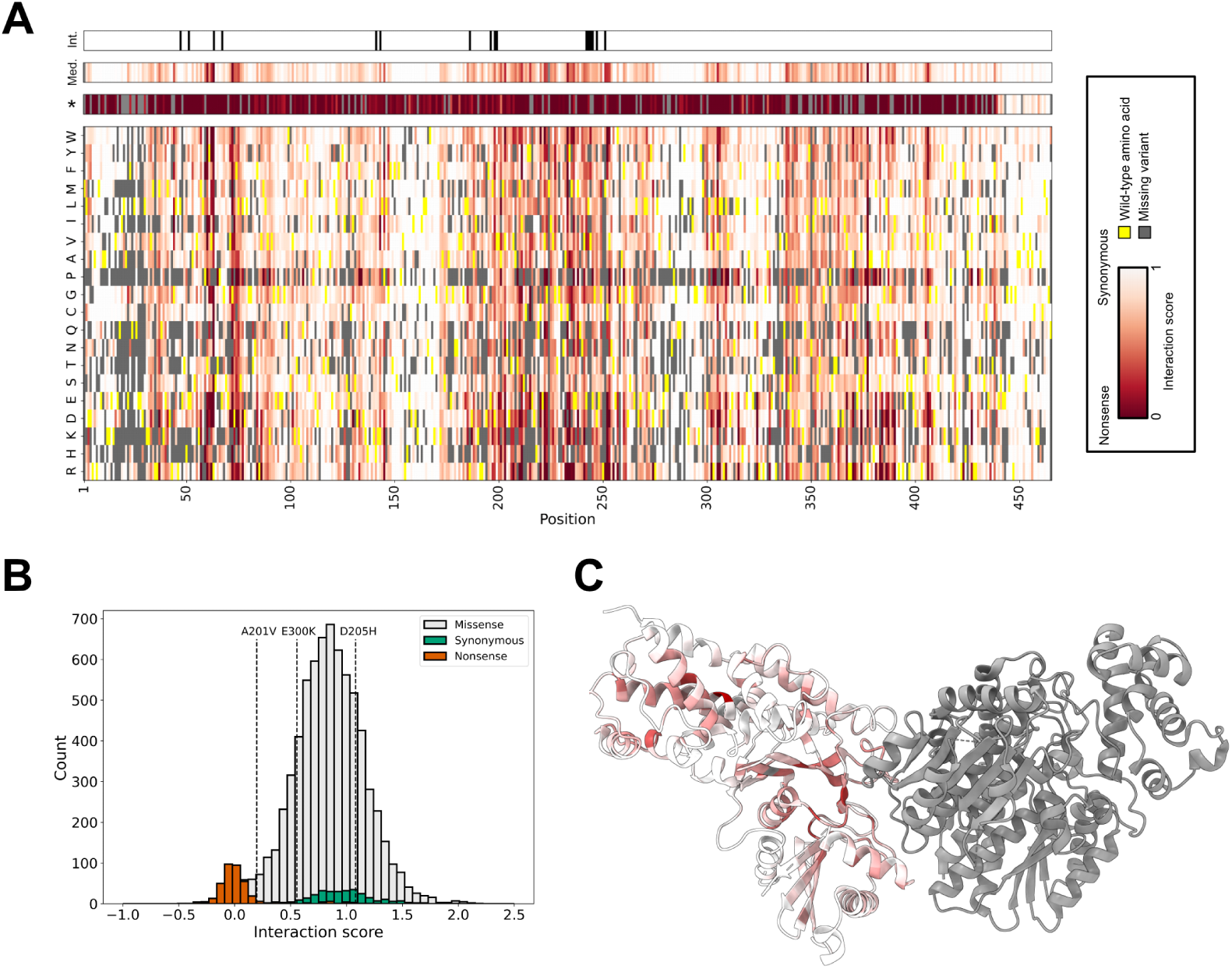
Map of GCK–GKRP interaction. (A) Heatmap showing the interaction score of each possible amino acid substitution (y-axis) along the GCK sequence (x-axis). Variants with no interaction are shown in red, while wild-type-like interaction is shown in white. The wild-type residue at each position is shown in yellow. Gray indicates missing variants. Two bars above the heatmap shows the median interaction score at each position (Med.) and the interaction score of nonsense variants (*). At the top, residues within 4 Å of GKRP in the crystal structure are shown as black lines. (B) Interaction score distributions of missense, synonymous and nonsense variants. The interaction scores of three variants tested in the small scale assay are shown as stippled lines. The uncapped interaction scores was used for this plot. (C) The median interaction score of each GCK residue mapped onto the structure of the GCK–GKRP complex. GKRP is shown in gray. The coloring scheme is the same as in panel A. PDB ID: 4LC9 [19].

When we examined all interaction scores as three distributions of missense, synonymous and nonsense variants (Fig. 2B), it was clear that synonymous variants had a wider distribution than nonsense variants. This is likely a result of the reverse selection, which means that wild-type-like variants will deplete during selection, leading to fewer sequencing reads and higher uncertainty. In addition to synonymous variants, this greater uncertainty also applies to wild-type-like missense variants. The distributions of missense and synonymous variants both included variants with a score greater than one, which could either be due to increased interaction or uncertainties at higher scores due to reverse selection. We therefore tested six missense variants with a high interaction score (spanning 1.4 to 2) in the reverse two-hybrid assay, and none showed increased interaction (Fig. S1). Hence, interaction scores above one likely result from noise, and we therefore capped the interaction scores at one in both the heatmap (Fig. 2A) and in all following figures, unless otherwise indicated.

To further examine the interaction scores, we mapped the median interaction score at each position of GCK onto the structure of the GCK–GKRP complex (Fig. 2C) [19]. As expected, mutations at the interaction interface decreased interaction. There were, however, many residues outside the interaction interface where variants also decreased interaction. These residues appeared in most of the GCK structure, but concentrated at some buried residues in the large domain. Variants at these residues likely decreased protein abundance and thereby interaction, as seen for the initial test variants (Fig. 1B and C).

### Correlations between experimental data and clinical genetics

As seen above, the interaction scores not only captured interaction-specific effects. Therefore, we examined which aspects of GCK function that the interaction scores capture. To this end we compared interaction scores with various measures of GCK–GKRP interaction, GCK activity and clinical data.

First, we examined the interaction scores of 11 missense variants that were previously shown to perturb GCK–GKRP interaction [25] (Fig. 3A). These variants all showed low interaction scores with the highest-scoring variant (T60A) having an interaction score of 0.43. The interaction scores therefore capture variants that specifically perturb GCK–GKRP interaction.

**Figure 3.**
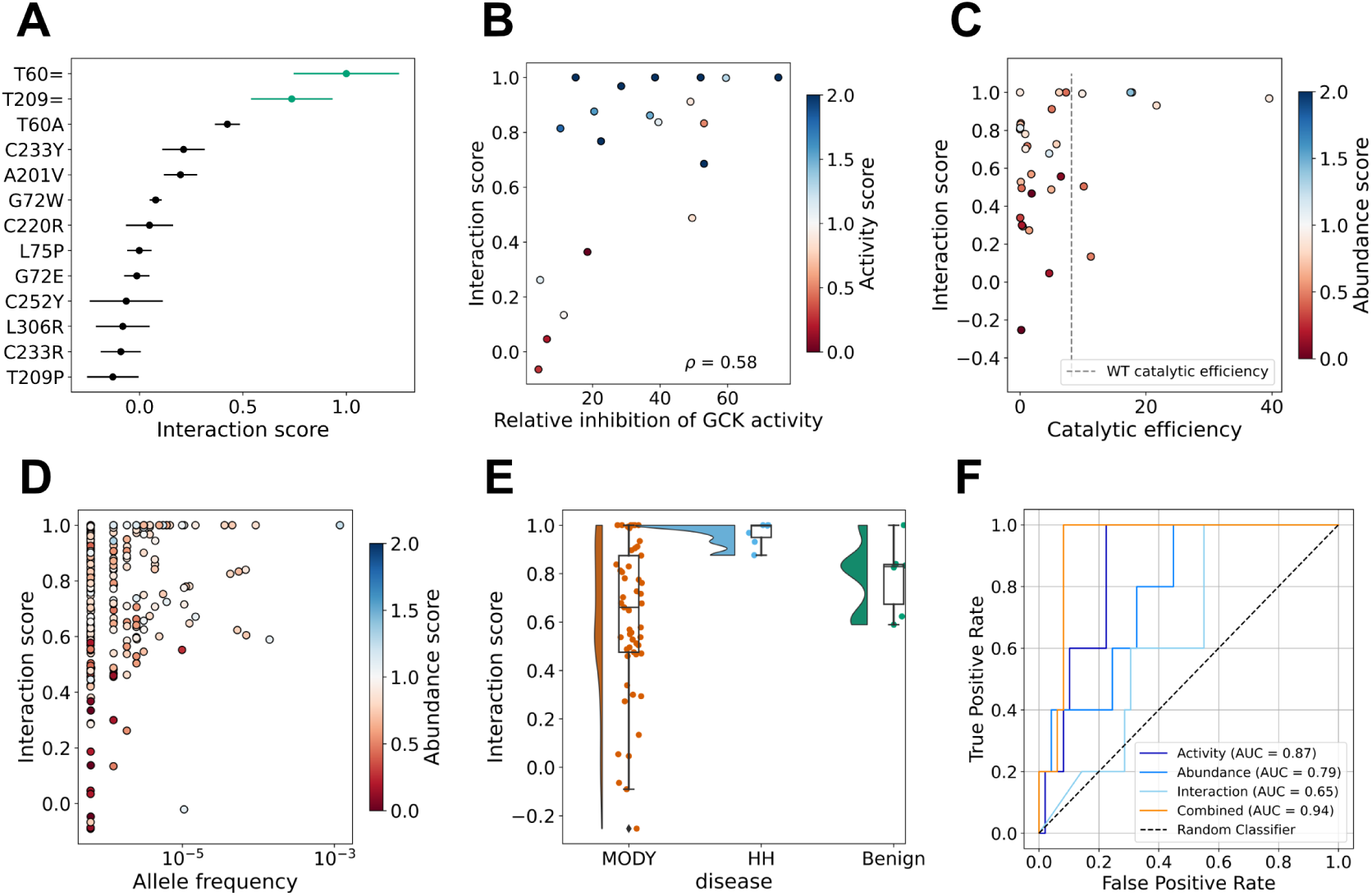
Comparing interaction scores with prior experimental and clinical data. (A) The interaction scores of 11 missense variants previously shown to decrease interaction [25]. The two synonymous variants at the top are included as references. (B) Correlation between interaction scores and GKRP-mediated GCK inhibition previously measured *in vitro* [29]. Variants are colored by their activity score [11], where red shows decreased activity, white shows wild-type-like activity and blue shows increased activity. (C) The interaction scores and catalytic efficiencies of 32 missense variants. The variants are colored by their abundance score [11], where red shows decreased abundance, white shows wild-type-like abundance and blue shows increased abundance. The catalytic efficiency of wild-type GCK is shown as a stippled line. The catalytic efficiencies were previously measured *in vitro* [10]. (D) The interaction scores and allele frequencies of missense variants found in gnomAD [30]. Variants are colored by abundance scores [12] with the same coloring scheme as in panel C. (E) Raincloud plot showing the interaction scores of 53 variants linked to GCK-MODY [31], 7 variants linked to HH [31] and 6 likely benign/benign (LB/B) variants [26]. (F) Receiver operating characteristic (ROC) curves showing the performance of activity scores, abundance scores, interaction scores, and the three scores combined on classifying GCK-MODY and LB/B variants.

To further examine the interaction-specific effects, captured by the interaction scores, we used a previously published dataset measuring GKRP-mediated GCK inhibition of 20 missense variants *in vitro* [29] (Fig. 3B). Interaction scores showed a reasonable correlation with relative GCK inhibition (Spearman’s *ρ*: 0.58). In addition, most of the variants that were not inhibited by GKRP *in vitro* but did interact with GKRP in our assay, displayed increased activity. Therefore, the deviation between interaction scores and *in vitro* inhibition might partially be explained by hyperactive variants that are able to bind to GKRP but are not inhibited. In conclusion, although the two datasets measure slightly different aspects of the GCK–GKRP interaction, the interaction scores partially capture inhibition measured *in vitro*.

Next, we examined whether there was a connection between a variant’s interaction score and *in vitro* catalytic efficiency (Fig. 3C) [10]. Variants with a high catalytic efficiency showed wild-type-like interaction, while variants with wild-type-like or lower catalytic efficiency showed interaction scores spanning from wild-type-like to complete loss of interaction. Variants decreasing both catalytic efficiency and interaction also decreased GCK protein abundance. Therefore, variants that decrease activity do not appear to decrease interaction specifically, but instead might structurally perturb GCK leading to decreased protein abundance.

Having compared interaction scores with different experimental measures of GCK function, we next examined the potential clinical relevance of these scores, noting that variants in pancreatic GCK are also relevant to liver GCK, as both isoforms are derived from the same gene, and that GCK displays distinct but overlapping functions in liver and pancreas [32]. First, we compared interaction scores with GCK allele frequencies from gnomAD (Fig. 3D) [30]. Variants with higher allele frequencies had interaction scores spanning from around 0.5 to 1, while variants with lower allele frequencies had interaction scores spanning from complete loss of interaction (0) to being wild-type like (1). Most of the variants with a low interaction score and low allele frequency also decreased protein abundance (Fig. 3D) and/or activity (Fig. S2A). There were, however, seven variants with a low allele frequency (<10^−5^), a wild-type-like activity score (>0.66 and <1.18), and a wild-type like abundance score (> 0.58) that all had interaction scores below 0.5. Of these seven variants, five were present in ClinVar, and of these ClinVar entries, three (K143N, P145L, V338M) were listed as variants of uncertain significance (VUS) and two (T209M, S263P) as likely pathogenic/pathogenic (LP/P). The interaction scores therefore appear to capture two disease-causing variants that are not identified by activity or abundance scores. Potentially, these variants affect interaction specifically. However, whether perturbing interaction with GKRP alone is enough for a variant to be disease-causing is unknown, and at least might result in a milder phenotype as the GCK–GKRP interaction is only relevant in hepatocytes and not in pancreatic *β*-cells.

To further examine any potential clinical relevance of interaction scores, we looked at the scores of 6 likely benign/benign (LB/B) variants [26], 7 variants linked to HH [31] and 53 variants linked to GCK-MODY [31] (Fig. 3E). Variants linked to HH did not affect interaction, while the benign variants varied a bit more in interaction scores, with the lowest-scoring variant, D217N, having an interaction score of 0.59. Finally, variants linked to GCK-MODY spanned the entire range of interaction scores. However, the GCK-MODY variants that decreased interaction also decreased GCK protein abundance (Fig. S2B). Again, this suggests that the interaction scores provide limited clinical value beyond variant abundance.

To examine whether the interaction scores provided any further clinical value when considering activity and abundance scores, we analyzed the ability of GCK scores to classify variants linked to GCK-MODY (Fig. 3F). We note that this analysis is limited by the low number of known LB/B variants (10 in ClinVar) with all three scores (5). To examine the ability of combined scores to classify variants, we defined the combined score as the minimum of the scores combined for each variant. Given the variants available, combining all three scores was the best classifier (AUC=0.94) (Fig. 3F), with activity and abundance scores combined performing similarly (AUC=0.93) (Fig. S2C). Finally, combined activity and interaction scores performed only slightly better (AUC=0.90) than activity scores alone (AUC=0.87) (Fig. S2D). Hence, for the GCK-MODY variants analyzed, interaction scores provide limited additional clinical value and mainly report on variant abundance, which, as expected, is better captured by abundance scores. The interaction scores may however still provide clinical value for pathogenic GCK variants not linked to GCK-MODY, although this remains to be examined.

In addition, when looking at the three score sets individually, the activity scores are the best classifier (AUC=0.87) as expected (Fig. 3F), while abundance scores are slightly worse (AUC=0.79) and interaction scores worse still (AUC=0.65). This suggests, as expected, that activity scores provide the most clinical value, as all pathogenic variants are expected to perturb activity directly or indirectly, while many of these, but not all, will decrease protein abundance [12, 33].

In conclusion, the interactions scores partly reflect the ability of GCK variants to interact with GKRP, but they also to a large extent reflect variant abundance.

### Correlations and deviations between interaction and abundance scores

Given that the interactions scores partly reflect protein abundance, we next examined the relationship between interaction and abundance scores. We compared the median interaction and abundance scores along the GCK sequence (Fig. 4A). The two medians generally showed the same pattern, suggesting that part of the interaction score reflected protein abundance, as expected. The two median scores, however, deviated in two aspects.

**Figure 4.**
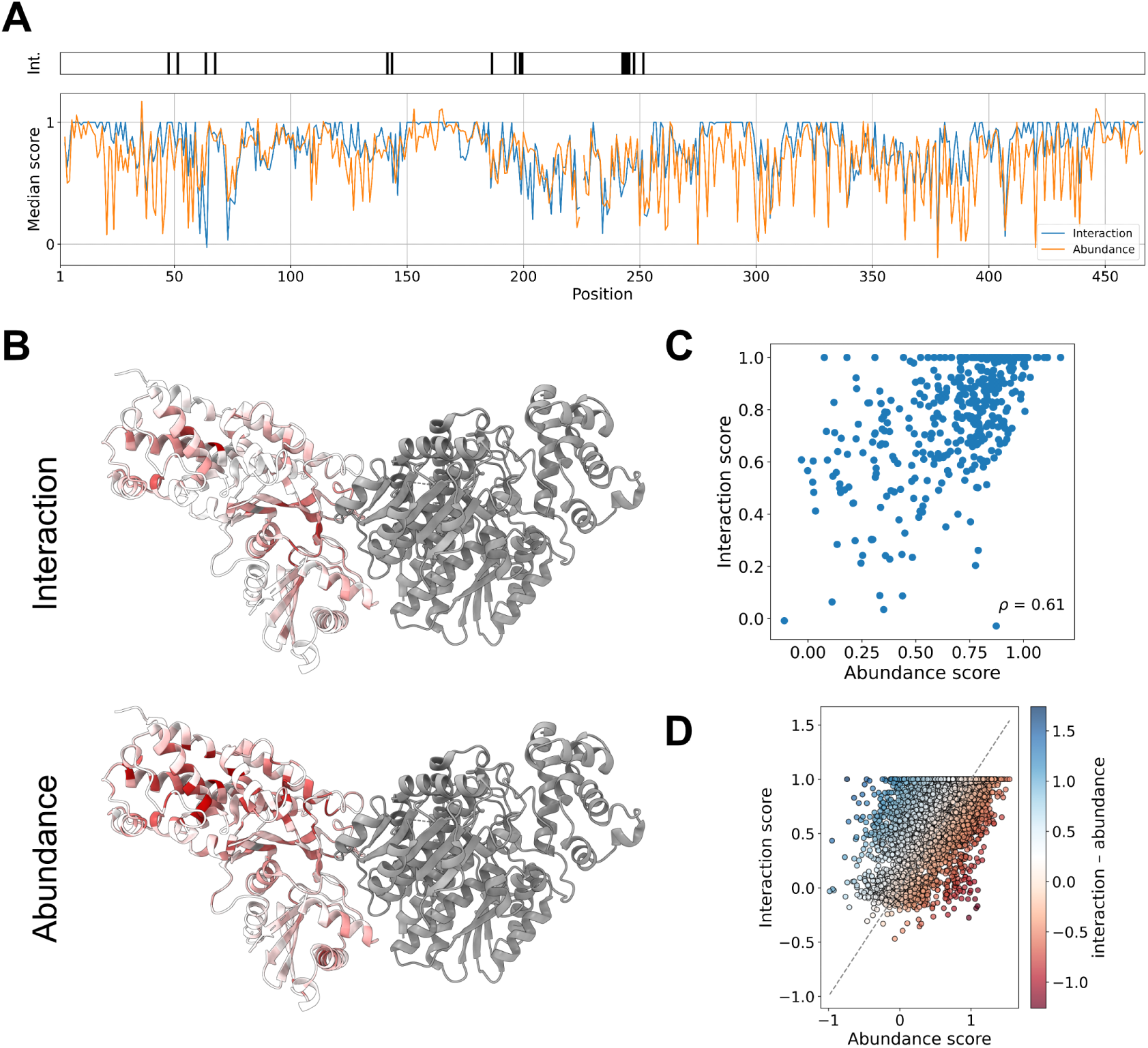
Variant effects on GCK interaction and abundance. (A) Plot of median interaction and abundance scores for each position along the GCK sequence. At the top, residues within 4 Å of GKRP in the crystal structure are shown as black lines. (B) Median interaction and abundance scores mapped onto the structure of the GCK–GKRP complex. Red shows nonsense-like interaction/abundance and white shows wild-type-like interaction/abundance. GKRP is shown in gray. PDB ID: 4LC [19]. (C) Plot showing the correlation between median interaction and abundance scores. (D) The correlation between variant-level interaction and abundance scores. Each variant is colored by the delta score calculated by subtracting abundance scores from interaction scores. Red shows variants that decrease interaction most, white shows variants that affect interaction and abundance equally and blue shows variants that decrease abundance most. A stippled line shows a perfect correlation between the two scores.

First, the median abundance score was generally lower, compared to the interaction score (Fig. 4A). When mapping the two median scores onto the structure of the GCK–GKRP complex (Fig. 4B), it was similarly clear that mutations were less detrimental for interaction. This was especially evident for buried residues in the large domain. Although the median interaction and abundance scores showed a reasonably high correlation (Spearman’s *ρ*: 0.61, Fig. 4C), many residues had a lower median abundance score. One possible explanation for this tendency is that unstable variants might be stabilized by binding to GKRP in the interaction assay, leading to higher protein abundance and thereby a higher interaction score compared with the abundance score.

Second, the two medians also deviated in that there were small regions where the median interaction score was lower than abundance such as around residues 60-70 and 200 (Fig. 4A). Some of these regions were found at the interaction interface of the the GCK–GKRP complex (Fig. 4B). Although it was only a few residues where variants decreased interaction more than abundance (Fig. 4C), many of these residues were not in the direct vicinity of GKRP in the structure of the complex (Fig. S3A, within 4 Å of GKRP). This suggest that variants in residues outside the interaction interface can also perturb interaction specifically. The mechanistic basis for this is, however, unknown, but could reflect allosteric effects.

To look further into interaction–abundance correlations and deviations, we examined the variant-level correlation between interaction and abundance scores, which was not perfect (Spearman’s *ρ*: 0.52) (Fig. S3B). We noticed a surprising pattern where low-abundance variants clustered in two populations with interaction scores around 0.5 and 0. This clustering was caused by nonsense variants scoring differently than missense variants in the two assays (Fig. S3C). For abundance scores, nonsense variants behaved similarly to low-abundance missense variants. In contrast, for interaction scores, nonsense variants generally showed lower interaction scores than low-abundance missense variants. Presumably, most nonsense variants result in a truncated GCK protein with an incomplete interaction interface, and will therefore show complete loss of interaction. In contrast, although missense variants may perturb the GCK structure, GKRP may in many cases still be able to bind GCK, and this interaction could potentially stabilize an unstable GCK protein variant.

Collectively, the above results show that the interaction scores partially reflect GCK variant abundance, but also that deviations between the two scores might identify variants that (1) perturb interaction specifically and (2) can be stabilized by binding to GKRP. To look further into these two groups of variants, we subtracted abundance scores from interaction scores to generate a delta score (Fig. 4D). A negative delta score suggests that a variant affects interaction most and therefore might identify the interaction-specific variants. A positive delta score suggests that a variant affects abundance most, which could potentially indicate that the variant is stabilized by GKRP in the interaction assay.

### Identifying GCK variants that affect GKRP interaction

Having calculated the delta scores, we examined what characterized variants with neutral, negative or positive delta. First, positive-delta variants generally had lower activity scores than neutral- or negative-delta variants (Fig. 5A). Second, both positive- and negative-delta variants generally had a higher weighted contact number (WCN) compared to neutral-delta variants (Fig. 5B) (the WCN for the GCK super-open state as previously calculated [12] was used). This suggests that variants decreasing either interaction or abundance more than the other are generally more buried than variants affecting interaction and abundance equally. Third, positive-delta variants were generally predicted to destabilize the GCK protein more than neutral- or negative-delta variants (Fig. 5C) (ΔΔ*G* calculated for the super-open state previously [12]). Collectively, these data suggest that positive-delta variants decrease activity, destabilize GCK, and are found in buried residues. Potentially, these variants decrease activity through loss of abundance, and their abundance may be increased by binding to GKRP.

**Figure 5.**
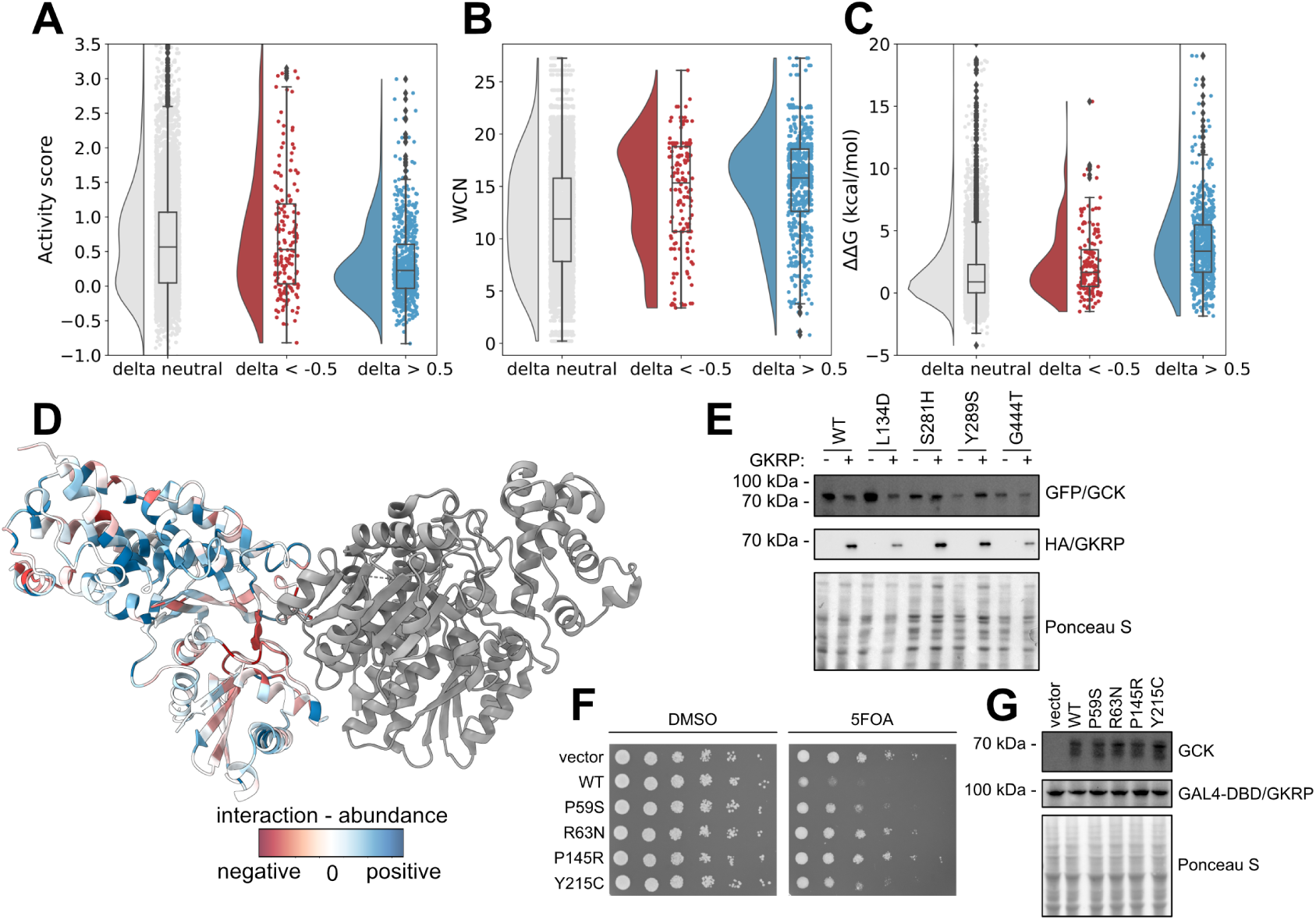
Delta scores identify variants stabilized by GKRP and variants that perturb interaction. (A) Raincloud plot showing the activity score distributions for variants grouped by their delta score. (B) Raincloud plot showing the weighted contact number (WCN) distributions for variants grouped by their delta score. (C) Raincloud plot showing the distributions of ΔΔ*G* values (calculated using the super-open GCK conformation) for variants grouped by their delta score. (D) The structure of the GCK–GKRP complex colored by median delta score. Red shows residues where variants decrease interaction most, white shows residues where variants affect interaction and abundance equally and blue shows residues where variants decrease abundance most. PDB ID: 4LC9 [19]. (E) Western blot showing the protein levels of GCK and GKRP in the *hxk*1Δ*hxk*2Δ*glk*1Δ yeast strain co-expressing EGFP-GCK (wild type or the shown variants) and a vector control or HA-GKRP. (F) Reverse two-hybrid growth assay using yeast co-expressing GAL4-DBD-GKRP and GAL4-AD-GCK (wild type or the shown variants). DMSO is used as a control while 5FOA selects for variants with decreased interaction. (G) Western blot showing the GCK and GKRP protein levels in the reverse two-hybrid setup.

To further examine the delta scores, we looked at the median across the GCK sequence. Both residues with a positive or negative median delta were spread throughout the GCK sequence (Fig. S4), although they still clustered together in small regions. When we mapped the median delta score onto the structure of the GCK–GKRP complex, residues with a negative median delta clustered at the interaction interface, as expected (Fig. 5D). There were, however, also residues outside the interaction interface where variants specifically perturbed interaction. In general, residues with a positive median delta clustered at buried residues in the large domain (Fig. 5D). As also seen above, this suggests that variants in buried residues in the large domain destabilize GCK, leading to decreased protein abundance that may be rescued by GKRP interaction.

We used low-throughput experiments on individual variants to examine whether positive- and negative-delta variants were indeed stabilized by GKRP and interaction-specific, respectively. First, we tested whether four positive-delta variants (L134D, S281H, Y289S, G444T) were stabilized by GKRP. We co-expressed wild-type GCK and the four missense variants with either a vector control or GKRP (Fig. 5E). Unexpectedly, we saw that wild-type GCK showed reduced protein levels when GKRP was co-expressed. Potentially this could be an indirect effect caused by the high expression level of GKRP (samples were diluted 1:20 for the GKRP blot relative to GCK). The same was seen for the L134D variant. For the three remaining variants, however, their protein levels were unchanged or increased by GKRP co-expression (Fig. 5E). This suggests that, relative to wild-type GCK, three (S281H, Y289S, G444T) of the four tested variants are stabilized by GKRP. Consequently, positive delta values can relatively well pinpoint variants stabilized by GKRP.

Lastly, we tested whether four negative-delta variants (P59S, R63N, P145R, Y215C) specifically perturbed interaction. In the reverse two-hybrid assay, all four variants showed increased growth and thereby decreased interaction compared with wild-type GCK (Fig. 5F). Their protein levels, assessed by western blotting, were similar to wild-type GCK (Fig. 5G), showing that the four variants specifically decrease interaction. Collectively, these results support that negative delta values can identify variants that perturb GCK–GKRP interaction.

## Conclusions

Missense variants can affect a broad range of molecular phenotypes, and multidimensional experiments can help disentangle the molecular origins of how missense variants perturb functions and may cause disease [34, 35]. In this study, we combined deep mutational scanning with reverse two-hybrid to examine how GCK single amino acid substitutions affect interaction with GKRP. We found that variants perturbing interaction were concentrated at the interaction interface and at buried residues in the large domain. Accordingly, many variants perturbing interaction were likely due to protein destabilization, and a major driver of loss of interaction therefore unsurprisingly appears to be loss of abundance.

To isolate variants that specifically perturbed interaction, and not protein abundance, we compared interaction scores with our previously obtained GCK abundance scores [12]. Variants specifically perturbing interaction were found at the interaction interface, as expected, but also at residues structurally far from GKRP in the GCK–GKRP complex. How these variants perturb the interaction is not known, but similar observations have previously been suggested to be evidence of allosteric effects on binding [36]. Importantly, the protein-wide identification of variants that perturb GCK–GKRP interaction provides information on this interaction in a cellular context, in contrast to a crystal structure. Our mapping of the interaction is, however, still potentially affected by using the pancreatic isoform of GCK, despite the physiologically relevant isoform for this interaction being the hepatic isoform. Still, the two isoforms only differ in the 15 N-terminal residues, which are structurally far from GKRP.

When comparing interaction and abundance scores, we also found that abundance scores were generally lower, suggesting that GKRP stabilized unstable GCK variants. We subtracted abundance scores from interaction scores to identify variants with a potential for restabilization. By stabilizing these low abundance variants, it may be possible to rescue GCK function. Although we show stabilization by GKRP, which also inhibits activity, these variants could potentially be targeted by e.g. pharmacological chaperones to rescue function. This may, however, be especially complicated for GCK due to the important balance between different conformations for appropriate activity, and this balance may be perturbed by preferential binding of small molecules to different conformations.

Collectively, we now have activity, abundance and interaction scores for 7128 GCK single amino acid substitutions, which have allowed us to gain a deep understanding of this important enzyme. The interaction scores together with our previous data on GCK variant abundance and activity, may inform on GCK variants perturbing only GCK–GKRP interaction with more subtle phenotypic effects than GCK variants linked to PNDM, HH or GCK-MODY. As seen for the common GKRP allele P446L, associated with raised triglycerides and reduced fasting blood glucose [22], changes in GCK–GKRP interaction can have phenotypic effects, although the extent of such effects for GCK variants in humans remains to be characterized.

## Methods

### Buffers

TE buffer: 10 mM Tris/HCl, 1 mM EDTA, pH 8.0. SDS sample buffer (4x): 250 mM Tris/HCl, 40% glycerol, 8% SDS, 0.05% pyronin G, 0.05% bromophenol blue, pH 6.8. SDS sample buffer was diluted to 1.5x in water and 2% *β*-mercaptoethanol was added before use. Wash buffer: 50 mM Tris/HCl, 150 mM NaCl, 0.01% Tween-20, pH 7.4. PBS: 6.5 mM Na_2_HPO_4_, 1.5 mM KH_2_PO_4_, 137 mM NaCl, 2.7 mM KCl, pH 7.4.

### Plasmids

The protein sequence of human pancreatic GCK (UniProt P35557-1) was codon optimized for yeast expression and cloned into pDONR221 (Genscript). The selected missense variants studied in low throughput were generated by Genscript. For reverse two-hybrid experiments, *GCK* was cloned into pDEST22 from the ProQuest Two-Hybrid System (Invitrogen) using Gateway cloning (Invitrogen). In this way, the N-terminus of GCK was fused to the GAL4 activation domain. To examine the abundance of GCK variants with and without co-expression of GKRP, *GCK* was cloned into pAG416GPD-EGFP-ccdB (Addgene plasmid 14316 ; http://n2t.net/addgene:14316 ; RRID:Addgene_14316) [37] using Gateway cloning (Invitrogen).

The protein sequence of human GKRP (UniProt Q14397-1) was codon optimized for yeast expression and cloned into pDONR221 (Genscript). For reverse two-hybrid experiments, *GKRP* was cloned into pDEST32 from the ProQuest Two-Hybrid System (Invitrogen) using Gateway cloning (Invitrogen). In this way, the N-terminus of GKRP was fused to the GAL4 DNA-binding domain. To examine the abundance of GCK variants in presence of GKRP, codon-optimized *GKRP* was cloned into pDONR221 with a 5’ HA tag (Genscript). *HA-GKRP* was then cloned into pAG415GPD-ccdB (Addgene plasmid 14146 ; https://www.addgene.org/14146/ ; RRID:Addgene_14146) [37] using Gateway cloning (Invitrogen).

### Yeast strains

For reverse two-hybrid experiments, GCK and GKRP were expressed in the MaV203 yeast strain from the ProQuest Two-Hybrid System (Invitrogen). To examine the abundance of GCK variants in absence and presence of GKRP, GCK variants (and GKRP) were expressed in the *hxk1*Δ *hxk2*Δ *glk1*Δ strain generated previously [11].

MaV203 yeast cells were cultured in Yeast extract-Peptone-Dextrose (YPD) medium (2% D-glucose, 2% tryptone, 1% yeast extract) prior to transformation, and in synthetic complete (SC) medium (2% D-glucose, 0.67% yeast nitrogen base without amino acids, 0.2% drop out (USBiological)) following transformation. *hxk1*Δ *hxk2*Δ *glk1*Δ yeast cells were cultured in YP and SC medium with D-galactose instead of D-glucose. Yeast cells were transformed using the LiAc/SS carrier DNA/PEG method [38].

### Reverse two-hybrid growth assay

To examine GCK-GKRP interaction using the reverse two-hybrid assay, yeast cells were grown overnight and harvested in the exponential phase (1200 g, 5 min, RT). Cells were washed in sterile water (1200 g, 5 min, RT) and resuspended in sterile water. All cell resuspensions were adjusted to an OD_600nm_ of 0.4 and were then used for a five-fold serial dilution in water. The dilution rows were spotted in drops of 5 µL onto agar plates without leucine and tryptophan containing 0.1% 5FOA (Thermo Scientific, R0812) dissolved in DMSO or, as a control, DMSO. The plates were briefly air dried and were incubated at 30 °C for two to three days.

### Protein extraction

Proteins were extracted from yeast cells using the NaOH method [39]. Yeast cells were grown overnight and, in exponential phase, 3 OD_600nm_ units were harvested in Eppendorf tubes (17,000 g, 1 min, RT). Cell pellets were shaken with 100 µL of 0.1 M NaOH (1400 rpm, 5 min, RT). Cells were then spun down (17,000 g, 1 min, RT) and the supernatant was removed. The pellets were dissolved in 100 µL 1.5x SDS sample buffer (1400 rpm, 5 min, RT), and were then boiled for 5 min before SDS-PAGE.

### Electrophoresis and blotting

To examine the protein levels of GCK variants, we used SDS-PAGE followed by western blotting. First, SDS-PAGE was used to separate proteins in yeast extracts by size on 12.5% acrylamide gels. Proteins were then transferred to 0.2 µm nitrocellulose membranes, and the membranes were subsequently blocked in 5% fat-free milk powder, 5 mM NaN_3_ and 0.1% Tween-20 in PBS. The blocked membranes were incubated with a primary antibody overnight at 4 °C. The following day, membranes were washed 3 times 10 minutes with Wash buffer, and were then incubated for 1 hour with a peroxidase-conjugated secondary antibody at RT. Then membranes were again washed 3 times 10 minutes in Wash buffer. The washed membranes were incubated for 3 minutes with ECL detection reagent (Amersham GE Healthcare), and were immediately developed using a ChemiDoc MP Imaging System (Bio-Rad). The primary antibodies were: anti-GCK (Abcam, ab88056), anti-GAL4 DBD (Abcam, ab135397), anti-GFP (Chromotek, 3H9 3h9-100), anti-HA (Roche, 11867423001). The secondary antibodies were: HRP-antirat (Invitrogen, 31470), HRP-antirabbit (Dako, P0217), HRP-anti-mouse (Dako, P0260).

### Glucokinase library

#### Cloning

Three regional GCK libraries were previously generated in pENTR221 [11]. The libraries span aa 2–171 (region 1), 172–337 (region 3), and 338–465 (region 3) of the GCK sequence. Each entry library was cloned into the pDEST22 destination vector using Gateway cloning. One large-scale Gateway LR reaction was mixed for each regional library: 199.1 ng pENTR221-GCK library, 450 ng pDEST22 vector, 6 µL Gateway LR Clonase II enzyme mix (ThermoFisher), TE buffer to 30 µL. LR reactions were incubated overnight at RT. Then, reactions were incubated with 3 µL proteinase K (37 °C, 10 min). Subsequently, 4 µL of each regional LR reaction was transformed into 100 µL NEB 10-beta electrocompetent *E. coli* cells using electroporation. Cells were recovered in NEB 10-beta outgrowth medium (37 °C, 1 hour), and were then plated on LB medium with ampicillin. Plates were incubated overnight at 37 °C. A minimum of 500,000 colonies were obtained for each transformation. Transformed cells were scraped from the plates using sterile water, and cells corresponding to 400 OD_600nm_ units were used for plasmid DNA extraction (Nucleobond Xtra Midiprep Kit, Macherey-Nagel).

#### Yeast transformation

The three regional pEXP22-GCK plasmid libraries were transformed into MaV203 yeast cells as described previously using the 30x scale-up [40]. Prior to library transformations, the MaV203 yeast strain was transformed with pEXP32-GKRP. MaV203 yeast cells expressing pEXP32-GKRP were grown overnight at 30 °C. The following day, the exponential culture was diluted with 30 °C SC-leucine medium to an OD_600nm_ of 0.3 in approximately 150 mL. The culture was then incubated at 30 °C with shaking for 5-6 hours. The cells were harvested, washed two times in sterile water (1200 g, 5 min, RT), and resuspended in a transformation mix (7.2 mL 50% PEG, 1.08 mL 1.0 M LiAc, 300 µL 10 mg/mL single-stranded carrier DNA, 30 µg plasmid library, sterile water to 10.8 mL). Cells were then heat-shocked in a 42 °C water bath for 40 minutes. During incubation, the cell suspension was mixed by inversion every 5 minutes. Following heat-shock, cells were harvested (3000 g, 5 min, RT) and the cell pellet was resuspended in 30 mL of sterile water. To determine the transformation efficiency, 5 µL of the resuspended cells were plated in duplicate on SC-leucine-tryptophan medium. The remaining cell suspension was diluted to an OD_600nm_ of 0.2 in SC-leucine-tryptophan medium, and was then incubated for two days at 30 °C.

If the library transformations resulted in a minimum of 500,000 transformants, two cell pellets of 9 OD_600nm_ units were harvested (17,000 g, 1 min, RT) as preselection samples (two technical replicates). The pellets were stored at -20 °C prior to DNA extraction.

In parallel to the library transformations, pEXP22-GCK wild type was transformed into the MaV203 yeast strain expressing pEXP32-GKRP using the small-scale transformation protocol [38].

#### Selection

To select for GCK-GKRP interaction, the three regional yeast libraries were grown in duplicate on medium containing 0.1% 5FOA. For each library and repeat, 3.6 OD_600nm_ units of transformed cells were harvested and washed three times with sterile water (1200 g, 5 min, RT). The cell pellet was resuspended in 500 µL sterile water and the cell suspension was plated on a BioAssay dish (245mm x 245 mm) with SC-leucine-tryptophan medium containing 0.1% 5FOA (Thermo Scientific, R0812). The plates were incubated at 30 °C for two days. Following selection, cells were collected in 30 mL sterile water by scraping. Samples post selection were collected by harvesting 9 OD_600nm_ units of the cells (17,000 g, 1 min, RT). The cell pellets were stored at -20 °C prior to DNA extraction.

In parallel, MaV203 yeast cells expressing pEXP32-GKRP and pEXP22-GCK wild-type were used for selection as above, except that for each wild-type replicate 0.4 OD_600nm_ units of cells were plated on a petri dish.

For the three regional libraries and the GCK wild-type control, two replicates of yeast cell pellets pre- and post-selection were used for plasmid DNA extraction. The ChargeSwitch Plasmid Yeast Mini Kit (Invitrogen) was used to extract plasmid DNA.

#### Sequencing

To determine the change in variant frequency following selection, we sequenced the regional libraries pre- and post-selection. The libraries were sequenced in short tiles to allow sequencing of both strands, thereby reducing the impact of base-calling errors. Each regional library was sequenced in the following tiles: region 1 (tile 1–5), region 2 (tile 6–10), and region 3 (tile 11–14).

First, each tile was amplified from the plasmid DNA extracted from yeast. The plasmid DNA from each regional replicate was adjusted to equal concentrations, and was then used as template in a PCR to amplify each tile, with each reaction consisting of: 20 µL Phusion High-Fidelity PCR Master Mix with HF Buffer (NEB), 1 µL 10 µM forward primer, 1 µL 10 µM reverse primer, 18 µL plasmid library template. Primer sequences are available in the supplementary data (SKG_tilenumber_fw/rev). Tiles were amplified using the following PCR program: 98 °C 30 sec, 21 cycles of 98 °C 10 sec, 63 °C 30 sec, 72 °C 60 sec, followed by 72 °C 7 min and 4 °C hold.

Second, tiling PCR products were used in an indexing PCR to allow for multiplexing. Each reaction consisted of the following: 20 µL Phusion High-Fidelity PCR Master Mix with HF Buffer (NEB), 2 µL 10 µM i5 indexing adapter, 2 µL 10 µM i7 indexing adapter, 1 µL 1:10 diluted PCR product, 15 µL nuclease-free water. Illumina index adaptors were added using the following PCR program: 98°C 30 sec, 7 cycles of 98 °C 15 sec, 65 °C 30 sec, 72 °C 120 sec, followed by 72 °C 7 min and hold at 4 °C.

Third, the amplicons were pooled, and 100 µL of this pool was run on a 4% E-gel EX Agarose Gel (Invitrogen), followed by gel extraction. The quality and size of the gel-extracted pooled amplicons were analyzed using a 2100 Bioanalyzer system (Agilent). Finally, the pool of amplicons was quantified using the Qubit dsDNA HS Assay Kit (Thermo Fisher Scientific) and the Qubit 2.0 Fluorometer (Invitrogen), prior to paired-end sequencing using an Illumina NextSeq 550.

## Data analysis

The TileSeqMave (https://github.com/jweile/tileseqMave, version 1.3.1) and TileSeq mutation count (https://github.com/RyogaLi/tileseq_mutcount, version 0.6.945) pipelines were used to calculate variant interaction scores from sequencing data.

## Availability of data and materials

The datasets supporting the conclusions of this article are available in Additional file 2.

Sequencing reads are available at the NCBI Gene Expression Omnibus (GEO) repository (accession number: GSE304851).

Interaction scores have been deposited in the MaveDB repository (accession number: urn:mavedb:00001 a).

The activity scores that were generated in a prior study are available at Zenodo (https://doi.org/10.5281/zenodo.7636310).

The abundance scores that were generated in a prior study are available at Zenodo (https://doi.org/10.5281/zenodo.10837280).

The code for generating the plots found in the paper is available at GitHub (https://github.com/KULL-Centre/_2025_Gersing_GCKinteraction) with an MIT license.

## Competing interests

K.L-L. holds stock options in and is a consultant for Peptone Ltd. All other authors declare no competing interests.

## Funding

The present work was funded by the Novo Nordisk Foundation (https://novonordiskfonden.dk) challenge program PRISM (to K.L.-L. and R.H.-P.), REPIN (to R.H.-P.) and NNF21OC0071057 (to R.H.-P.), and the Danish Council for Independent Research (Det Frie Forskningsråd) (https://dff.dk) 10.46540/2032 00007B (to R.H.-P.). Computing resources were supported via a grant from the Carlsberg Foundation (https://www.carlsbergfondet.dk/) CF21-0392 (to K.L.-L.). The funders had no role in study design, data collection and analysis, decision to publish, or preparation of the manuscript.

## Authors’ contributions

S.G. and S.Ø.D. performed the experiments. S.G., K.L.-L. and R.H.-P. analyzed the data. S.G., K.L.-L. and R.H.-P. conceived the study. S.G. wrote the paper.

## Acknowledgements

The authors thank Jochen Weile and Roujia Li who developed the TileSeq pipeline used for analyzing sequencing data, Kristoffer E. Johansson for assistance with access to computing resources, Line Pedersen for assistance with Illumina sequencing, Sven Larsen-Ledet for help with the Bioanalyzer system, and Anne-Marie Lauridsen for technical assistance. We acknowledge access to the sequencing and core computing facilities at the Department of Biology, University of Copenhagen. Protein structures were visualized and rendered using UCSF ChimeraX (v1.4), developed by the Resource for Biocomputing, Visualization, and Informatics at the University of California, San Francisco [41, 42].

In Figure 1, panels A and D were created using BioRender.

## Supplementary information

Additional file 1: Supplementary figures. Additional file 2: Supplementary data.

