## Additional file 1: Supplementary figures for "Mapping the GCK–GKRP interaction landscape using deep mutational scanning by reverse two-hybrid screening"

### Supporting information for: Mapping the GCK–GKRP interaction landscape using deep mutational scanning and reverse two-hybrid

**Sarah Gersing**[1*], **Sofie Ø. Dahl**[1], **Kresten Lindorff-Larsen**[1], **Rasmus Hartmann-Petersen**[1*]

***For correspondence:**
 (S.G.);
 (R.H.-P.)

[1]The Linderstrøm-Lang Centre for Protein Science, Department of Biology, University of Copenhagen, Ole Maaløes Vej 5, DK-2200 Copenhagen, Denmark

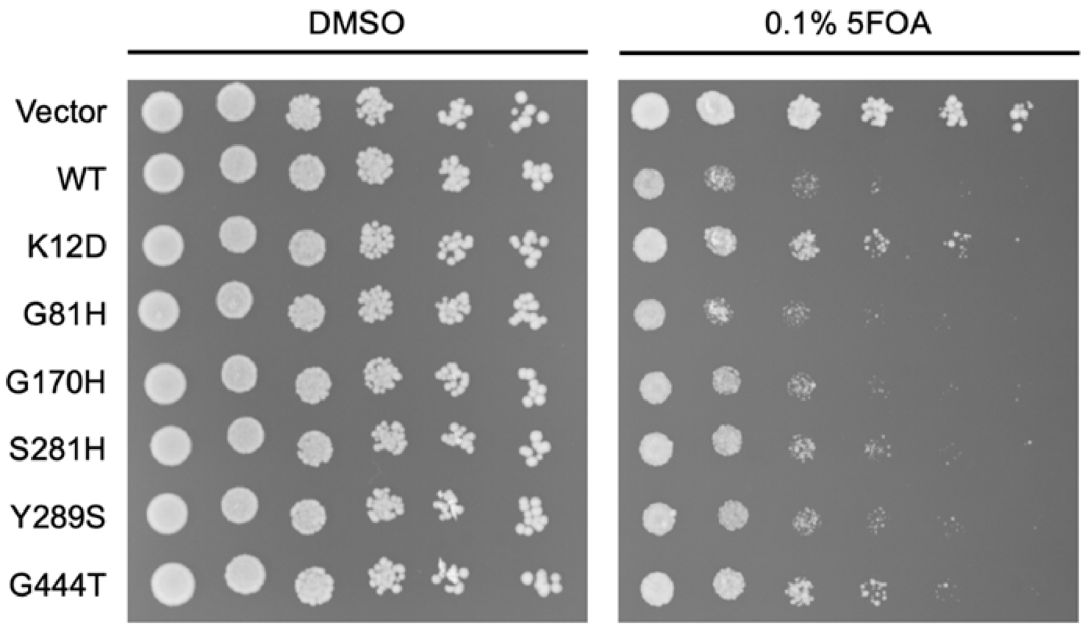

**Figure S1.** Reverse two-hybrid assay of variants with a high interaction score. Reverse two-hybrid using yeast co-expressing GAL4-DBD-GKRP and GAL4-AD-GCK (wild type and the shown variants). DMSO was used as a control while 5FOA selects for variants with decreased interaction. All the variants tested had an interaction score above 1, but did not show decreased growth compared to wild type, suggesting that the high interaction scores was caused by noise.

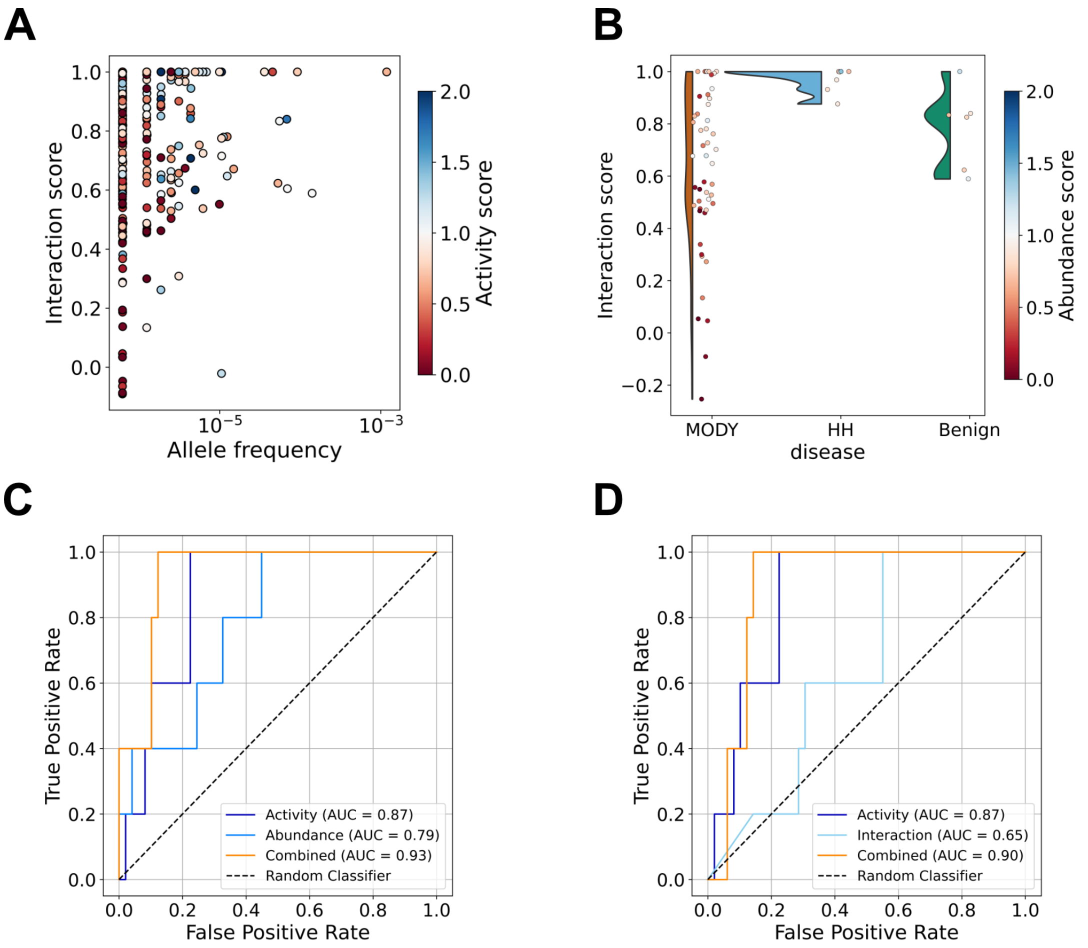

**Figure S2.** Examining the clinical value of interaction scores. (A) Plot showing the interaction scores and allele frequencies of GCK missense variants in gnomAD [1]. Variants are colored by their activity score. (B) Raincloud plot showing the interaction scores of 53 variants linked to GCK-MODY [2], 7 variants linked to HH [2] and 6 likely benign/benign (LB/B) variants [3]. Variants are colord by their abundance score. (C) Receiver operating characteristic (ROC) curves showing the performance of activity scores, abundance scores and the two scores combined on classifying GCK-MODY and likely benign/benign variants. (D) Receiver operating characteristic (ROC) curves showing the performance of activity scores, interaction scores and the two scores combined on classifying GCK-MODY and likely benign/benign variants.

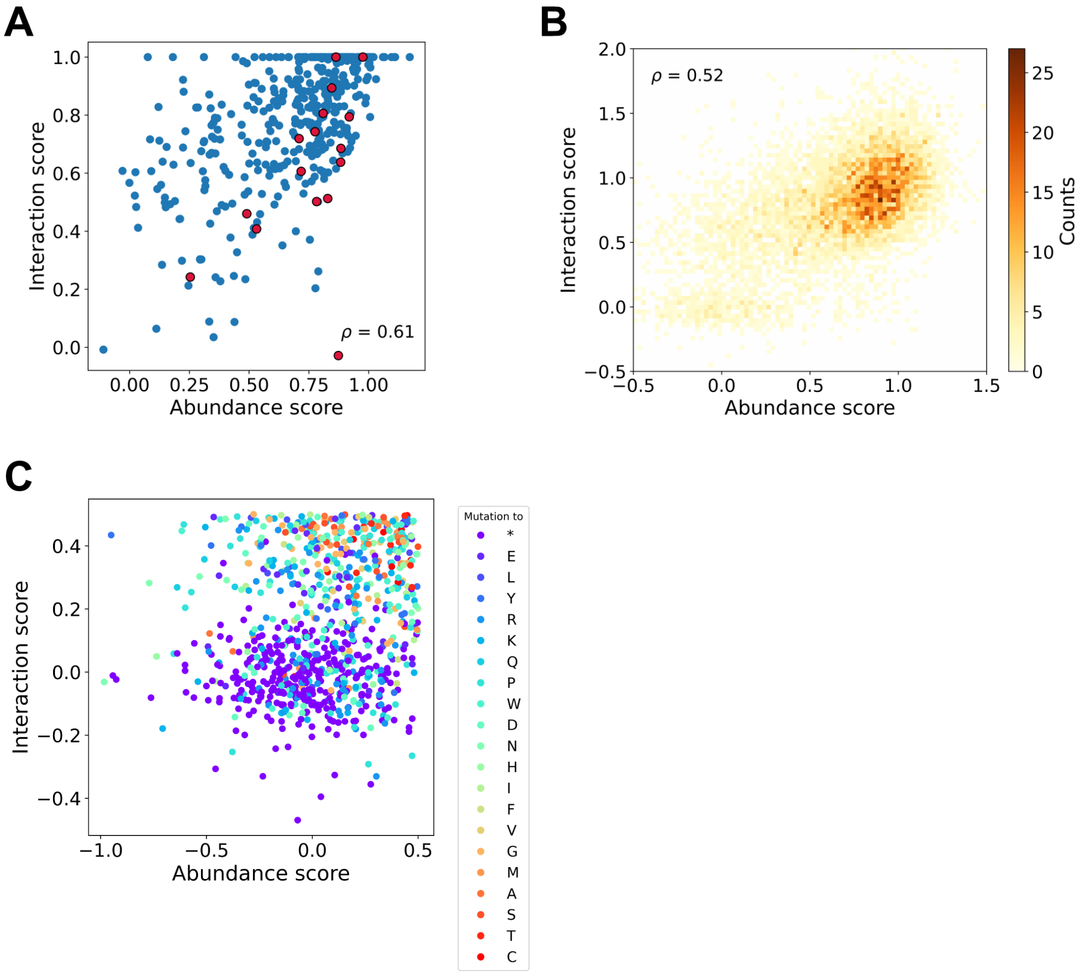

**Figure S3.** Comparing interaction and abundance scores. (A) Plot showing the correlation between median interaction and abundance scores. GCK residues within 4 Å of GKRP in the GCK–GKRP complex are colored red. PDB ID: 4LC9 [4]. (B) Plot showing the correlation between variant-level interaction and abundance scores. In this plot, uncapped interaction scores were used. (C) Plot showing that variants with an abundance score below 0.5 cluster in two groups in regard to their interaction score. This clustering is caused by the different interaction scores of missense and nonsense variants. Each variant is colored by the residue that they are mutated to.

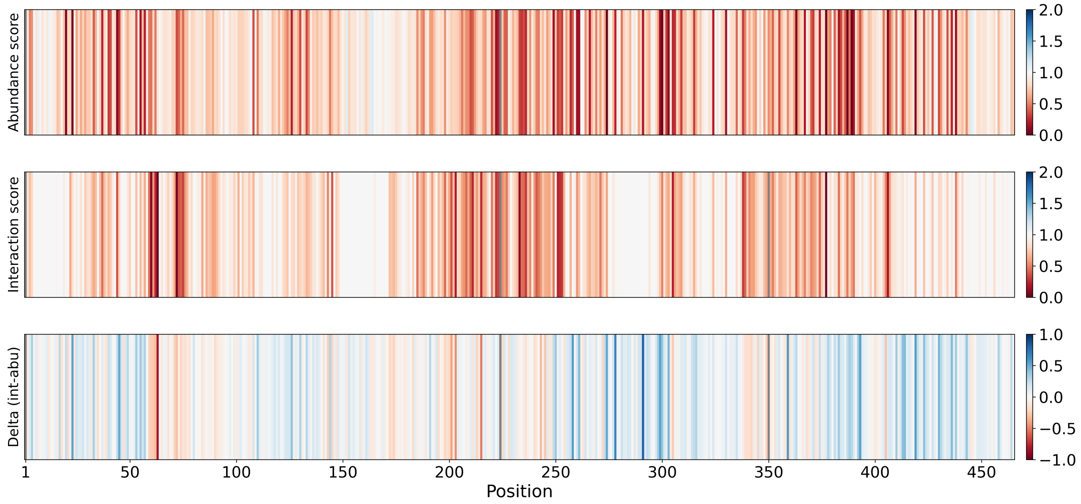

**Figure S4.** Median abundance, interaction and delta scores. Barcode plots showing the median abundance, interaction and delta score at each position along the GCK sequence. For abundance and interaction scores, red shows positions where variants decrease abundance/interaction and white shows positions where variants on average do not affect interaction/abundance. For delta scores, red shows positions where variants decrease interaction most, white shows positions where variants affect interaction and abundance equally and blue shows positions where variants decreased aundance most.
